# A subcellular spatial proteome of the non-photosynthetic euglenid *Euglena longa*

**DOI:** 10.64898/2026.09.28.755152

**Authors:** Michael J. Hammond, Yu-Ping Poh, Anzhelika Butenko, Julius Lukeš, Jeremy G. Wideman

## Abstract

Euglena longa is a heterotrophic model euglenid, of significant research interest as a unicellular eukaryote. To provide broad insight into the localisation of its proteome we used subcellular spatial proteomics (SSP), based on a label-free version of LOPIT-DC (Localisation of Organelle Proteins by Isotope Tagging after Differential Centrifugation), in which organelles are separated by differential centrifugation and each protein is assigned to a compartment from its abundance profile across the fractions. We detected 5,909 proteins, using protein abundance profiles across fractions to resolve 16 different subcellular compartments, such as the plastid, pellicle, and peroxisomes, and assigned high-confidence localisations to 2,955 of them. This study serves as a proteomic dataset to resolve protein compartmentalisation for this euglenid, enabling evolutionary, biological and functional insight and analysis. We are drafting manuscripts concerning the metabolic nature of the plastid, the evolutionary origins of the pellicle, and the functions of the peroxisome. Investigators interested in using this data prior to publication are encouraged to contact us to avoid overlap and to facilitate coordinated and collaborative use of this resource.

## Introduction

Euglena longa is a unicellular heterotrophic euglenid within the euglenozoan lineage of the protist supergroup Discoba. Euglenids are of broad research interest due to their biotechnological applications for biofuel and nutraceutical production, in addition to their ecological distribution and evolutionary position within the eukaryotic tree of life (Ginger, et al. 2024). Within euglenozoans, euglenids branch sister to the glycomonads, comprising diplonemids, which are significant inhabitants of marine plankton communities (Tashyreva, et al. 2022), and kinetoplastids, a lineage of heterotrophic flagellates that includes various mammalian parasites (e.g., Leishmania and Trypanosoma) (Gull 2001). These three lineages of Euglenozoa share common morphological features like the microtubule corset, an organised network of microtubules surrounding the periphery of the cell, and a cellular cavity known as the flagellar pocket, containing flagella furnished with a paraflagellar lattice (Kostygov, et al. 2021). While both diplonemids and kinetoplastids have compartmentalised certain glycolytic enzymes to peroxisomes, termed ‘glycosomes’ (Andrade-Alviárez, et al. 2022), such localisations are not found in euglenid peroxisomes, rendering them as the comparative ‘outgroup’ to this evolutionary trajectory.

The defining morphological trait (aka synapomorphy) of euglenids is their pellicle, a series of striated proteinaceous strips between the plasma membrane and microtubule corset (Cavalier-Smith 2017). A clade of the euglenid tree acquired photosynthetic plastids via a secondary endosymbiosis of green algae, enabling phototrophic lifestyles in addition to their varied heterotrophic living strategies (Ginger, et al. 2024). The most famous euglenid, Euglena gracilis, is an example of a photosynthetic euglenid. Euglena longa, though closely related, has lost its ability to photosynthesize, though it retains a colourless plastid and subsists as a secondary osmotroph. Outstanding questions remain over the capacity of its non-photosynthetic organelle, and whether notable metabolic repurposing or specialisation has occurred, particularly in light of its indispensability in contrast to the plastid of Euglena gracilis (Hadariová, et al. 2017).

Efforts to sequence high-quality euglenid genomes have been hampered by their large sizes (predicted 1-9 Gbp) and an abundance of repetitive regions (Fields, et al. 2025). The transcriptome assembly of E. longa predicts 55,704 proteins (Záhonová, et al. 2018), compared with over 36,000 from the transcriptome of E. gracilis (Ebenezer, et al. 2019), though the expression capacity of this entire proteome is unclear due to low proteomic recovery. Furthermore, most of these proteins are ‘hypothetical’, lacking any characterisation in euglenids or other eukaryotes. Thus, any localisation data is valuable to begin unravelling their functions.

To address this issue, we performed SSP using a modified LOPIT-DC fractionation workflow (Geladaki, et al. 2019), exploiting the recent success of SSP in resolving cellular compartments across a variety of other protists (Barylyuk, et al. 2020; Zítek, et al. 2022; Guérin, et al. 2023; Moloney, et al. 2023; Chen, et al. 2025; Hammond, et al. 2025; Jirsová, et al. 2025; Chisholm, et al. 2026; Hammond, et al. 2026). We ultimately identified 5,909 E. longa proteins via mass spectrometry and confidently assign localizations for nearly 3,000 of them to 16 different subcellular compartments.

## Results and Discussion

### Subcellular Fractionation and Proteomic Profiling of Euglena longa

Axenically grown cells were lysed and underwent fractionation by differential ultracentrifugation, before analysis via mass spectrometry. Briefly, ∼2.5 × 10^8^ axenically grown cells per quadruplicate were gently sonicated on ice to disrupt the integrity for the majority of cells while preserving internal organelles. The resulting lysate was separated by differential ultracentrifugation into 11 distinct fractions, with pellets harvested for downstream mass spectrometry analysis, with final supernatant representing fraction 11 (Table 1).

**Table 1:**
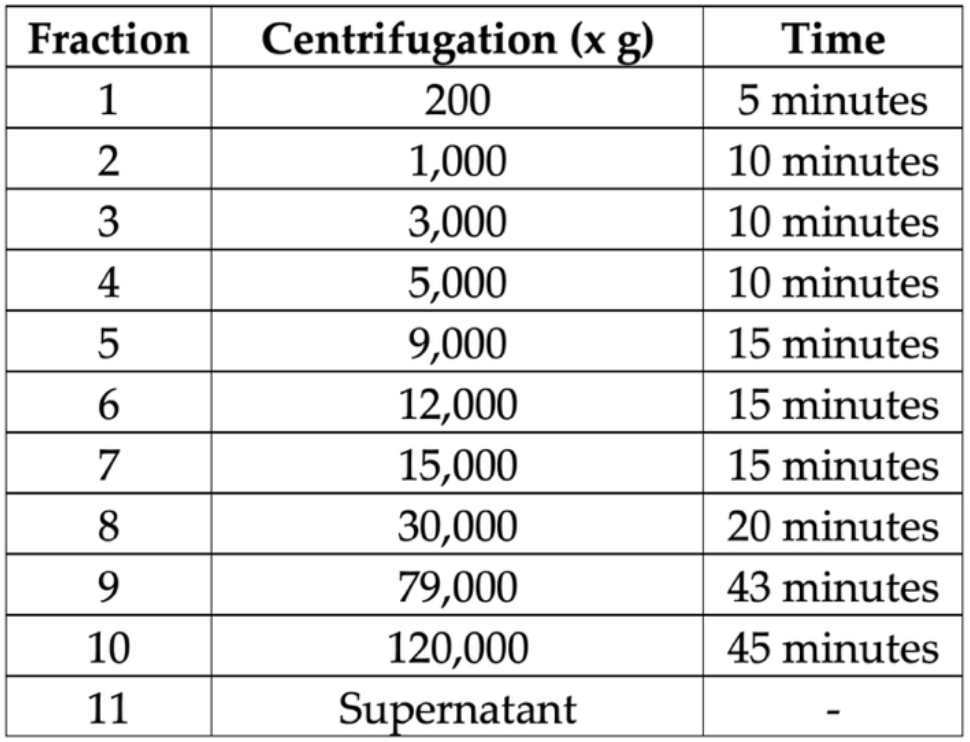
Centrifuge conditions employed for fractionation of E. longa cell lysate, based on Geladaki, et al. 2019.

Distinct enrichment of major organelles was confirmed by Western blotting using antibodies against established marker proteins for the mitochondrion (ATP synthase β) and nucleus (H3 histone) (Figure S1). Clear resolution of these markers across different fractions validated compartment separation, enabling subsequent proteomic analysis.

Label-free mass spectrometry analysis of the 11 fractions, performed in quadruplicate, identified 5,909 predicted proteins of the Euglena longa proteome. Protein abundances were normalised and missing values were imputed through neighbour averaging and zero methods, with enrichment profiles across combined fractions generated for each protein detected by mass spectrometry (Figure 1). Principal component analysis (PCA) was performed for dimensionality reduction, and the resulting principal components were subsequently visualised using t-distributed stochastic neighbour embedding (t-SNE). The t-SNE embedding revealed cluster-like patterns of proteins (Figure 1A), which were visualised according to their protein enrichment profiles (Figure 1B).

**Figure 1:**
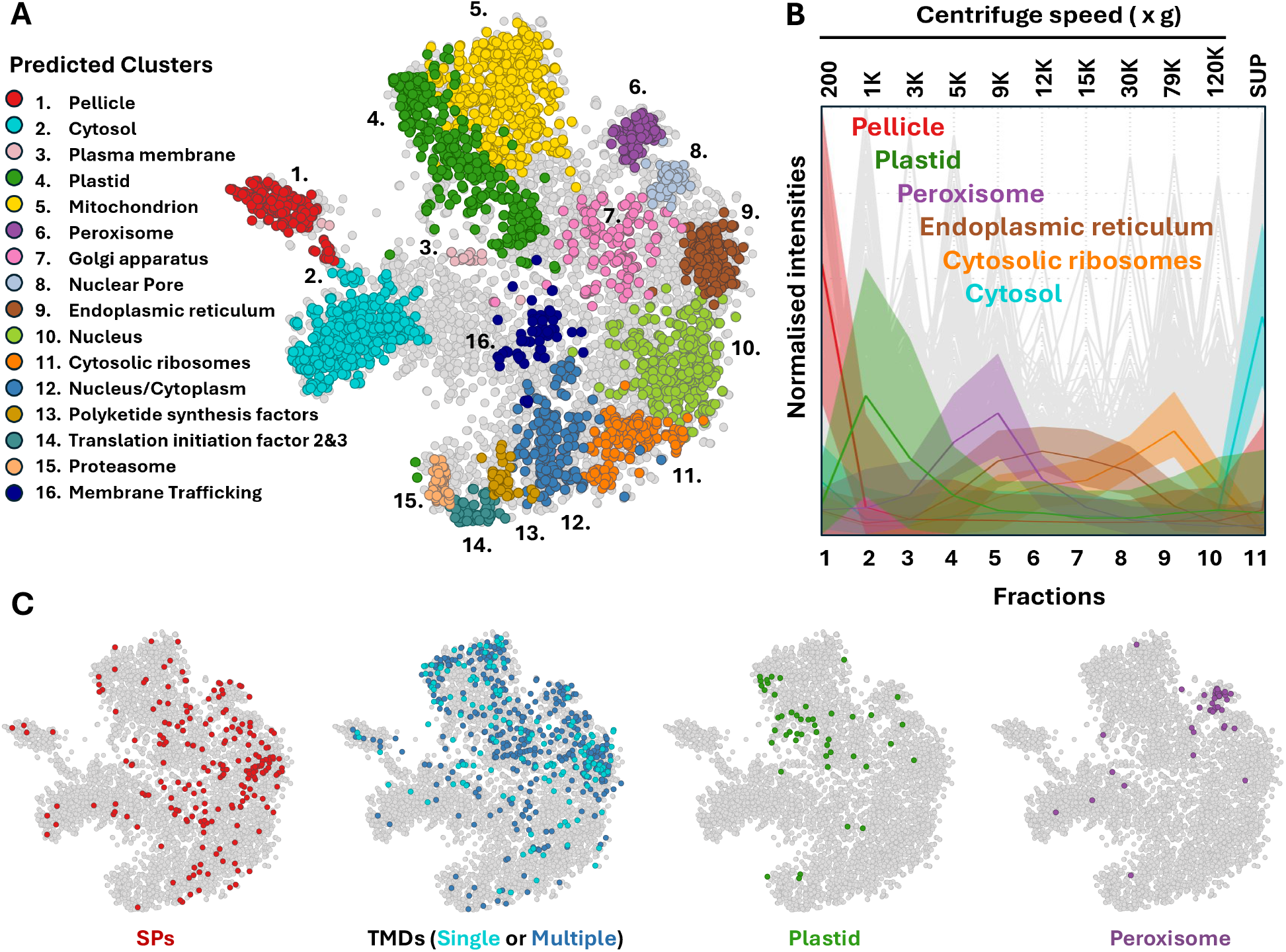
Subcellular proteome of Euglena longa. **A.** Protein localisations represented via t-SNE, predicted through supervised support vector machine clustering algorithm, trained on 14-20 marker proteins per organelle or compartment. **B.** Normalised abundance profiles were generated through mass spectrometry of collected fractions, with example compartments showing distinct fractional enrichment across one experimental quadruplicate. **C.** Subcellular localization prediction programs identified a variety of target peptide and domains across the subcellular plot, including signal peptides (SPs), Transmembrane domains (TMDs), plastid and peroxisome targeting sequences, corresponding to predicted clusters.

### Machine Learning-Based Assignment of Subcellular Localisation

To define subcellular compartments, we compiled a training set of 278 marker proteins (Table S1) with either experimental localization evidence within Euglena (Hammond, et al. 2020; Vanclová, et al. 2020) or those with strong homology to validated markers in other eukaryotes, particularly drawing on established proteomics for euglenozoan sister clades (Moloney, et al. 2023; Pyrih, et al. 2023; Hammond, et al. 2025). This curated set was used to train a support vector machine (SVM) classifier on these combined fractionation profiles. Application of the trained SVM with a median probability cutoff assigned subcellular localizations to 2,955 proteins (Figure 1A and Table S2), thereby providing the first predicted localisation for thousands of previously unannotated Euglena proteins.

The reliability of the SVM assignments was assessed using sequence-based prediction tools (Figure 1C). Predicted signal peptides were predominantly associated with the endoplasmic reticulum, Golgi apparatus, membrane trafficking pathway and plasma membrane (Teufel, et al. 2022). Transmembrane domains were enriched in all membrane-bound organelle clusters but predominantly absent in the nuclear and cytosolic clusters, as expected (Hallgren, et al. 2022). We additionally noted an enrichment of single transmembrane domain containing proteins within the endoplasmic reticulum. Prediction of targeting signals to organelles additionally shows correlation for regions of the plastid and peroxisome (Odum, et al. 2024). Furthermore, unsupervised k-means clustering with k = 16, matching the number of marker-defined compartments, shows broad correspondence with the SVM assignments (Figure S2), supporting the accuracy of fractional-based predictions, and the conclusions that can thus be drawn from this dataset.

## Conclusions

Here we report the first subcellular proteome of Euglena longa, which provides an investigative resource to explore protein function, compartmental organization, and cellular architecture within euglenids.

We aim to use this proteome to clarify what metabolic functions the now-colourless plastid still performs within this euglenid, and whether functional specialisations or new pathways via enzyme targeting has accompanied this organelle alteration. The well-resolved pellicle cluster provides us with the opportunity to explore how this peripheral structure has evolved following the divergence of euglenids from their sister-clades of diplonemids and kinetoplastids. Furthermore, we hope to characterise various sulphur-related enzymes and their subcellular localisations within the peroxisomes, as suggested by this dataset. Investigators interested in using these data are encouraged to contact the authors prior to publication to avoid overlap and to facilitate coordinated and collaborative use of this resource.

## Data availability

The mass spectrometry proteomics data have been deposited to the ProteomeXchange Consortium via the PRIDE partner repository with the dataset identifier PXD083867.

## Acknowledgements

J.G.W. was supported by a grant from the National Science Foundation (DBI-2119963) and a grant from the Gordon and Betty Moore Foundation (GBMF10600). J.L was supported by the Czech Grant Agency (23-06479X and 25-15298S).

## Methods

### Culturing and Cell Harvesting

Euglena longa strain CCAP 1204-17a, generously supplied by the Hampl lab from the BIOCEV institute, was cultivated axenically at 22 °C at 150 RPM within Cramers-Myers media, supplemented with ethanol to 0.8% of final volume. Late-stage logarithmic growth cultures were harvested by centrifugation at 1,500 x g for 5 minutes and washed twice in DF lysis buffer (0.25 M sucrose, 10 mM HEPES pH 7.4, 2 mM EDTA and 2 mM magnesium acetate). Cells were ultimately resuspended in 8ml lysis buffer for subsequent procedures.

### Cell Lysis and Fractionation

Cells were lysed by sonication (Heat Systems ultrasonics Serial number 5951 Model H-I) performed on ice with 15 second bursts at 60% power, repeated three times, interspersed with two-minute breaks. Cell lysate underwent fractional separation according to a differential centrifugation protocol as listed in Table 1 and described by Geladaki, et al. (2019).

### Western BloIing

Western blots were performed to assess compartment distribution to confirm that subcellular proteins were successfully separated into distinct fractions.

Pellets obtained from differential centrifugation were resuspended in 2X Laemmli buffer (4% SDS, 20% glycerol, and 120 mM Tris-HCl pH 6.8) at 95 °C until pellets dissolved. Resuspended protein fractions were quantified using Pierce™ BCA Protein Assay Kit (Thermo Scientific) according to the manufacturer’s instructions. 10 µg of proteins supplemented with bromophenol blue were loaded onto Invitrogen™ Bolt™ Bis-Tris Plus Mini Protein Gels (4–12%, 1.0 mm) alongside the Full Range Amersham™ ECL™ Rainbow™ Marker (MilliporeSigma). Electrophoresis was performed at 135 V for 1h. Proteins were transferred to PVDF membranes using the iBlot™ 2 Transfer Stacks (Invitrogen™) and the iBlot™ 2 Western Blot Transfer Device following the default program optimized for Bolt™ gels (25V for 6 minutes). Membranes were activated with methanol and briefly rinsed with deionized water and blocked for 1 h at room temperature in Kroger™ 5% dry milk prepared in 1 × Tris-buffered saline (TBS) with gentle shaking. Primary antibody incubation was performed using rabbit-derived antibodies: anti-Mitochondrial ATP synthase β (supplied by Zíkova lab, polyclonal (Šubrtová, et al. 2015)), and anti-Histone H3 (Novus Biological, NB500-171, polyclonal).

Primary antibodies were diluted in blocking buffer (1:10,000 for ATP Sβ and 1:500 for histone H3), incubated for 1 h at room temperature with gentle rocking, and then overnight at 4 °C. Following three 10-min washes in TBS, membranes were incubated for 1 h at room temperature with Goat anti-Rabbit IgG (H&L) HRP-conjugated secondary antibody (ImmunoReagents Inc.; 1:5,000 in blocking buffer), washed three times for 10 min each in TBS-T (TBS + 0.05% Tween-20), and briefly rinsed in water. Protein bands were detected using the Pierce™ ECL Western Blotting Substrate (Thermo Scientific) and chemiluminescent signals were captured using an Azure 600 Biosystems imaging system.

### Sample preparation and LC–MS Analysis

Native protein pellets obtained from differential centrifugation were digested and desalted following the protocol for the S-Trap Micro Column (ProtiFi, USA). Protein concentration was quantified using the BCA assay (Thermo Fisher Sci.), while peptide concentration was measured using a fluorometric kit (Thermo Fisher Sci.).

### Liquid-chromatography tandem mass spectrometry

LC–MS/MS analyses were performed at the Biosciences Mass Spectrometry Core Facility, Arizona State University. Data-dependent mass spectra were collected in positive mode using an Orbitrap Fusion Lumos mass spectrometer coupled with an UltiMate 3000 UHPLC (Thermo Fisher Sci.). Peptides were fractionated on an Easy-Spray LC column (50 cm × 75 µm ID, PepMap C18, 2 µm, 100 Å) with an upstream trap column. Each sample was analyzed in technical triplicate. LC–MS settings: electrospray potential 1.6 kV, ion transfer tube temperature 300°C, and the “Universal” peptide analysis method. Full MS scans (375–1,500 m/z) were acquired at a resolution of 120,000 with 3 s cycles. The RF lens was set to 30%, AGC to “Standard,” and monoisotopic peak determination included charge states 2–7. Dynamic exclusion was 60 s with a 10 ppm mass tolerance. MS/MS spectra were acquired in centroid mode with a quadrupole isolation window of 1.6 m/z and CID energy of 35%. Peptides were eluted over a 240-min gradient at 0.25 µL/min using 2%–80% acetonitrile/water: 0–3 min (2%), 3–75 min (2%–15%), 75–180 min (15%–30%), 180– 220 min (30%–35%), 220–225 min (35%–80%), 225–240 min (80%–85%).

### Raw data processing and quantification

The LFQ analysis was performed using Proteome Discoverer 2.4 (Thermo Fisher Sci.) based on the composite database: E. longa’s predicted proteome, and mitochondrial open reading frames (ORFs), raw files were searched with SequestHT using Trypsin as the enzyme, allowing up to three missed cleavages. Peptide length was set to 6–144 AAs, with precursor ion mass tolerance at 20 ppm, fragment mass tolerance at 0.5 Da, and a minimum of one peptide identified. Carbamidomethyl (C) was a fixed modification, while Acetyl (N-terminus), Met-loss (N-terminus), and oxidation of Met were dynamic modifications. A target/decoy strategy and 1.0% FDR were calculated using Percolator. Data were imported into Proteome Discoverer 2.4, and features were detected using the Minora Feature Detector algorithm. The area under the curve for aligned ion chromatograms was calculated to determine relative abundances.

### Raw data processing

Individual protein abundance profiles from all quadruplicates were merged to generate a comprehensive protein abundance dataset spanning all fractions. Proteins that were completely absent in any experiment were excluded from further analysis. The remaining proteins and their corresponding abundance values were imported into R and converted into an MSnSet object using the Bioconductor packages MSnbase (v2.24.2; Gatto and Lilley 2012) and pRoloc (v1.38.2; Gatto, et al. 2014). Missing values were then imputed by nearest-neighbour averaging followed by zero-filling, and the imputed dataset was used for downstream PCA. The resulting principal components were then visualized using t-Distributed Stochastic Neighbor Embedding (t-SNE) to present the similarities of normalised protein distribution across subcellular fractions (van der Maaten and Hinton 2008).

### Supervised Classification

SVM classification was performed on post-imputation marker protein abundance profiles using the Bioconductor pRoloc package (v1.38.2) in R. A curated set of 278 marker proteins (14-20 per subcellular compartment, across 16 compartments) was used to train the classifier. The optimal SVM parameters determined from the marker set were then applied to all proteins in the dataset, generating SVM scores ranging from 0 to 1, with 1 corresponding to marker protein confidence. Proteins without labels (non-marker proteins) were classified using these parameters, with weights applied according to the marker classes. Proteins with SVM scores below the global median were reset to “unknown” while proteins above the median were considered predicted to their corresponding compartments.

### Targeting sequence identification

SignalP 6.0 was first applied to identify signal peptides indicative of Sec/SPI-mediated targeting (Teufel, et al. 2022), employing proteins with a score greater than 0.8 as genuine. DeepTMHMM v1.0 (Hallgren, et al. 2022) was then used to predict transmembrane topology. DeepLoc 2.1 (Odum, et al. 2024) was further used as an alternative tool to predict targeting signals and putative localisations of all proteins.

**Figure S1:**
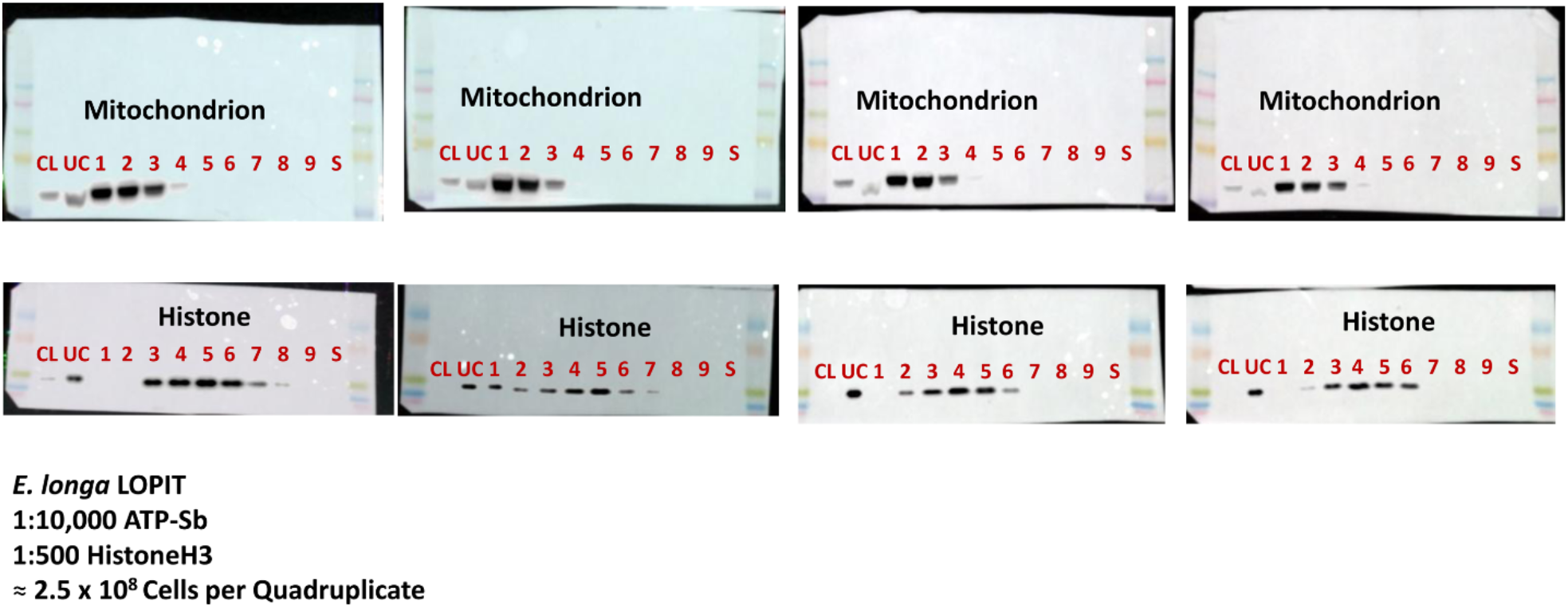
Western blot of protein distribution in quadruplicate fractions, including cell lysate (CL), and unlysed cells (UC) and supernatant (S). 10 µg of protein was loaded per lane.

**Figure S2:**
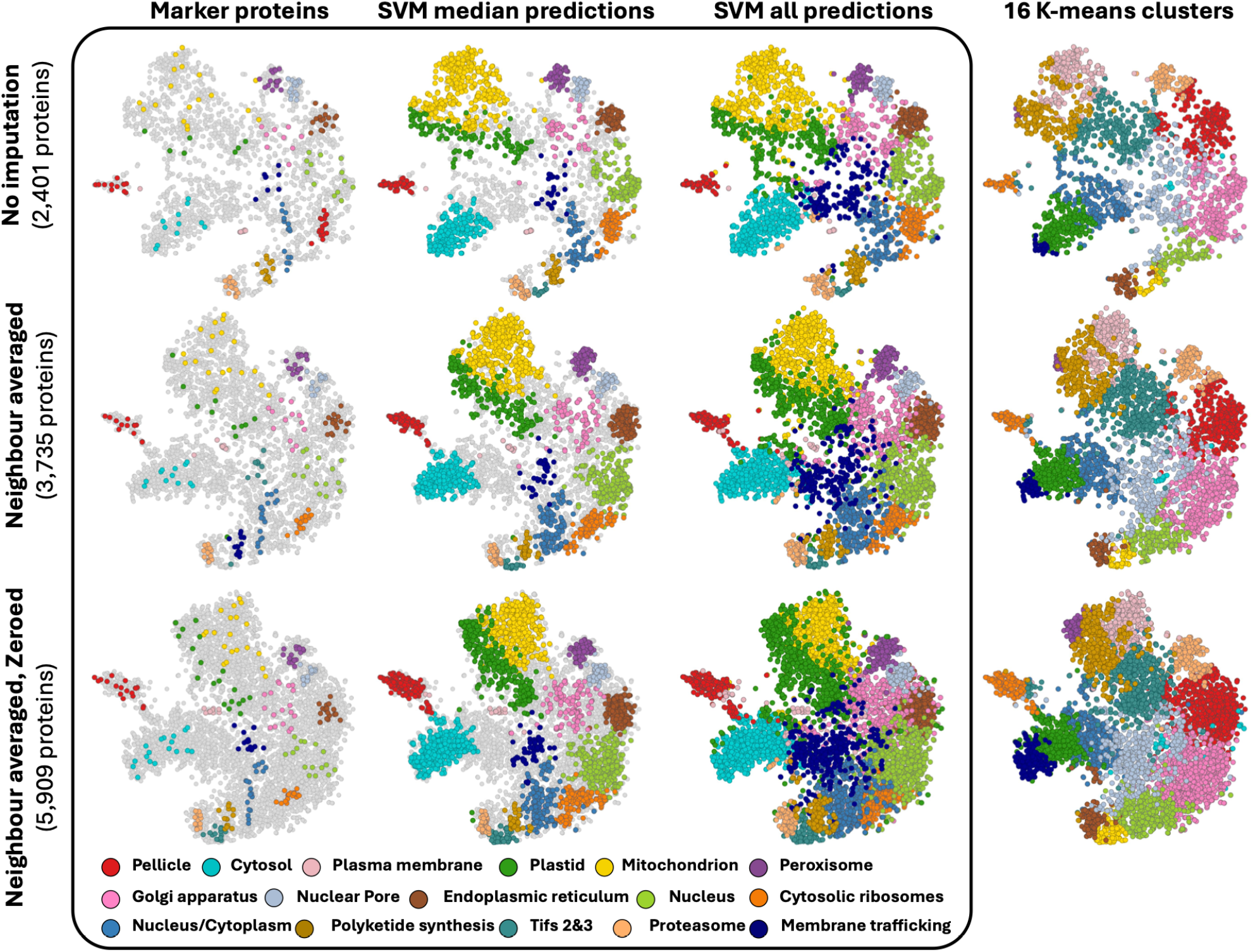
Euglena longa subcellular proteome, showing imputation conditions as well as marker proteins, SVM predictions and classifications below calculated median, along with unsupervised classification showing 16 K-means clusters.

**Table S1. Marker proteins used in this study.**

**Table S2. SVM scores for all proteins detected in this study.**

